# Sodium Pyruvate Ameliorates Influenza A Virus Infection *In Vivo*

**DOI:** 10.1101/2020.11.25.396978

**Authors:** Jessica M. Reel, Christopher R. Lupfer

**Author notes:** **Address correspondence to:** Dr. Christopher R. Lupfer, Department of Biology, Missouri State University, 901 S. National Ave., Springfield, Missouri, 65897.

## Abstract

Influenza A virus (IAV) causes seasonal epidemics annually and pandemics every few decades. Most antiviral treatments used for IAV are only effective if administered during the first 48 hours of infection and antiviral resistance is possible. Therapies that can be initiated later during IAV infection and that are less likely to elicit resistance will significantly improve treatment options. Pyruvate, a key metabolite, and end product of glycolysis, has been studied for many uses, including its anti-inflammatory capabilities. Sodium pyruvate was recently shown by us to decrease inflammasome activation during IAV infection. Here, we investigated sodium pyruvate’s effects on IAV *in vivo.* We found that nebulizing mice with sodium pyruvate decreased morbidity and weight loss during infection. Additionally, treated mice consumed more chow during infection indicating improved symptoms. There were notable improvements in pro-inflammatory cytokine production (IL-1β) and lower virus titers on days 7 post-infection in mice treated with sodium pyruvate compared to control animals. As pyruvate acts on the host immune response and metabolic pathways and not directly on the virus, our data demonstrate that sodium pyruvate is a promising treatment option that is safe, effective, and unlikely to elicit antiviral resistance.

## 1. Introduction

Influenza A virus (IAV) causes seasonal epidemics and periodic pandemics with significant morbidity and mortality. In the 2019-2020 flu season, the United States Center for Disease Control and Prevention (CDC) estimated 38 million IAV infections and 22,000 deaths. The most prevalent virus of the 2019-2020 season was the 2009 pandemic IAV (H1N1). Notably, during this season, was a higher rate of infections among children aged 0-4 and adults aged 18-49 years than in other recent seasons [1]. During pandemics, the emergence of novel viruses can cause severe complications with increased morbidity and mortality [2]. Due to the novelty of pandemic viruses, vaccines must be redesigned. Anti-viral therapies exist to treat IAV [3]. However, viral resistance to these therapies are always possible. Therefore, treatments that alter the host response to IAV infection and are less likely to result in evolution of resistance are desirable.

Studies have shown that IAV hijacks cellular metabolism to increase viral replication [4, 5]. Pyruvate (Pyr) (C_3_H_4_O_3_) is a central metabolite and key component in energy metabolism and cellular respiration. Pyr can enter directly into the mitochondria to produce ATP via the tricarboxylic acid cycle (TCA), which bypasses many of the metabolic regulatory pathways that control glycolysis [6, 7]. Mitochondrial oxidative phosphorylation is the most efficient way to produce ATP for cells. However, Pyr can also be used to make amino acids or be reduced to lactate via fermentation or the Warburg Effect [8, 9]. Reduction of Pyr is used to replenish NAD^+^ and increase uptake of necessary nutrients for rapidly dividing cells, such as immune and cancer cells [8–10]. The end goal is rapid proliferation, not energy efficiency, in most of these cases [10].

Pyr in its many forms (ethyl Pyr, pyruvic acid, pyruvate anion, sodium pyruvate, etc.) have been found to have many antioxidant-like benefits in several body’s systems. The molecule seems to be well tolerated in the body with little to no toxicity [11]. Additionally, Pyr has been found to have a plethora of beneficial effects on the cardiac system [12–14]. Moreover, increasing extracellular Pyr in the brain has been found to decrease neuron death during traumatic brain injury events [15, 16] and be protective to neurotoxic compounds [17]. When administered to mice of various ages, Pyr increased glycogen stores and brain energy metabolites, which could help with diseases such as Alzheimer’s [18]. Also, Pyr decreases epithelial permeability, inflammation, and bacterial translocation during intestinal ischemic reperfusion events [19, 20]. Bone and tissue inflammation models have shown that Pyr treatment lead to less destructive results of disease via anti-inflammatory properties [21, 22]. In relation to infectious disease, sodium pyruvate (NaPyr) (C_3_H_4_NaO_3_) can improve herpes simplex 2 virus infection *in vivo*, and our lab recently reported that NaPyr can regulate inflammation during IAV infection *in vitro* [23, 24].

In our previous study, we observed in mouse bone marrow derived macrophages (BMDM), that NaPyr has anti-inflammatory capabilities through altered metabolism [23]. The addition of NaPyr to BMDM decreased mitochondrial damage in response to IAV infection. These findings led us to further investigate NaPyr’s potential anti-viral and anti-inflammatory capabilities in a mouse model of IAV infection. Here we show that nebulizing NaPyr *in vivo* in WT C57BL/6J mice leads to decreased weight loss and increased chow intake over the course of IAV infection. Seven days p.i., animals treated with NaPyr displayed decreased pro-inflammatory cytokines (IL-1β) in the lungs and decreased virus replication.

## 2. Materials and Methods

### 2.1 Animal Welfare

WT C57BL/6J mice were bred and raised in the Temple Hall Vivarium at Missouri State University. Mice were euthanized via CO_2_ asphyxiation and cervical dislocation or cardiac puncture at humane end points or tissue collection. All breeding and experiment protocols were performed in accordance with Institutional Animal Care and Use Committee (IACUC) guidelines (protocols 19.005 and 19.019), the AVMA Guidelines on Euthanasia, NIH regulations (Guide for the Care and Use of Laboratory Animals), and the U.S. Animal Welfare Act of 1966.

### 2.2 Virus Production

The strain of IAV used in all experiments was influenza A/PR/8/34 H1N1 (PR8). PR8 stocks were generated by infecting pathogen-free hen’s eggs with 1000 PFU of PR8. Following a 3-day incubation, the allantoic fluid was harvested, centrifuged to remove debris, and stored at −80°C for later use.

### 2.3 *In Vivo* Infection and NaPyr Treatments

Mice were anesthetized on Day 0 via intraperitoneal injection of 80 mg/kg of Ketamine and 8 mg/kg of Xylazine. Mice were then infected intranasally with approximately 250 PFU of influenza A/PR/8/34 H1N1, diluted in 30 μL of phosphate buffered saline (PBS).

Mice used for subcutaneous (Sub-Q) injections of NaPyr were injected with 110mg/kg of body weight daily, divided into two doses, morning and evening. Mice that were treated with nebulized NaPyr were treated with Emphycorp’s clinical grade N115 (20mM NaPyr), or with 10 mM NaPyr (Fisher Bioreagents, BP356-100) diluted in PBS, or treated with PBS alone as control. Mice were treated three times a day for 20-minute per treatment. All mice were monitored for food/water availability and weighed daily for weight loss and/or becoming moribund. Mice were euthanized if experiment on Day 14, or day of sacrifice for tissue samples. Food intake was also monitored by weighing the food daily and averaging the change in food weight by the number of animals per cage.

### 2.4 Tissue Collection and Processing

Mice sacrificed for tissue samples on 7 p.i. were euthanized via CO_2_ asphyxiation and cardiac puncture as an adjunct. Lungs were taken from sacrificed mice for processing. Lungs were weighed and homogenized through a 70 μm cell strainer (Fisherbrand, 22363548) with a final volume of 4 mL of RPMI 1640 without serum or NaPyr (Hyclone, SH30027.01) per tissue sample. Samples were then centrifuged and aliquoted for future use.

### 2.5 Flow Cytometry for Innate and Adaptive Immune Cells

Lung homogenates were centrifuged at 400xg for 7 minutes to achieve cell pellet. After removal of the supernatant for other assays, red blood cells were lysed with ACK lysis buffer. Dead cells and debris were then removed by centrifugation in 37.5% Percoll (GE Healthcare, 17-0891-02) at 2000g for 20 minutes. Cells were then stained with fluorescent antibodies (**Table 1**). Samples were run on the Accuri C6 Flow Cytometer.

**Table 1:** Fluorescent antibodies used for FACS staining in preparation for flow cytometry. Antibodies were purchased from Tonbo or Biolegend.

| Fluorophore | Monocyte Stain | Cat# | Lymphocyte Stain | Cat# |
| --- | --- | --- | --- | --- |
| FITC | CD11c | 35-0114-U100 | CD4 | 35-0042-U100 |
| PE | Gr1 | 50-5931-U100 | CD8 | 100707 |
| PerCP 5.5 | CD3ε | 65-0031-U100 | CD3ε | 65-0031-U100 |
| APC | CD11b | 20-0112-U100 | CD19 | 115511 |

### 2.6 Viral Plaque Assay

The IAV plaque assay was performed using MDCK cells seeded at 2×10^5^ cells/well in 12-well plates in DMEM+ 5% FBS+ 1%Pen/Strep. 10-fold dilutions of the virus were prepared in RPMI 1640. MDCK cells were washed with PBS twice and 100μl of each virus dilution added to wells in 12-well plates and incubated at 37°C and 5% CO_2_ for one hour. Semisolid overlay was prepared as previously described [25]. TPCK-trypsin was added to a final concentration of 1.0 μg/ml. After a full hour of incubation, infection medium was removed from 12-well plates, 2ml of the warm overlay with TPCK trypsin was added to each well and allowed to solidify. Plates were turned upside down and incubated for 3 days at 37°C and 5% CO_2_. After incubation, the overlay was removed, and plaques counted after staining with 1% crystal violet in formalin.

### 2.7 Enzyme-Linked Immunosorbent Assay (ELISA)

Supernatant from homogenized lung tissue samples were analyzed for IL-1β and IL-6. ELISA kits were purchased from Ebioscience (88-7013-88, 88-7064-88) and assays performed according to the manufacturer’s recommendations. Plates were read at 450nm on a microplate reader (BioTek ELx808).

### 2.8 Statistical Analysis

Statistical analysis was performed using GraphPad PRISM9. For *In Vivo* weight loss and chow consumption during experiments, a two-way ANOVA was performed. For viral titer, cytokine and cell populations analysis, a Student’s t-test was performed. A p-value <0.05 was considered statistically significant.

## 3. Results

### 3.1 NaPyr is not toxic *In Vivo*

N115 is a clinical grade nasal spray containing 20nM NaPyr that has undergone safety and phase I, phase II and phase III clinical trials. The FDA is currently reviewing the administration of N115 for use in COPD patients with Idiopathic Pulmonary Fibrosis, or Idiopathic Pulmonary Fibrosis Patients alone (EmphyCorp, Cellular Sciences inc FDA submissions). Patient surveys indicated that use of N115 may decrease the incidence, symptoms and duration of respiratory infections too (**Table 2**). Millions of patients have been treated with N115 nasal spray in over 200 hospitals globally, which includes pregnant women, patients with allergic rhinitis, COPD patients, sinusitis, and patients with pulmonary fibrosis, with no adverse events reported. The use of the nasal spray in these patients demonstrates its safety and efficacy and the ability of NaPyr to reduce nasal congestion and inflammation. In a Phase III Placebo Controlled Clinical Trial with Idiopathic Pulmonary Fibrosis Patients, the N115 nasal spray demonstrated a statistically and clinically significant increase in nasal nitric oxide, FEV-1, SaO2, FVC, FEV-1/FVC ratios (52% to 86%). N115 reduced hypoxemia, and it also reduced lung inflammation, inflammatory cytokines, and coughing. Other studies confirm the safety of supplementation with NaPyr [11, 16, 26].

**Table 2.**
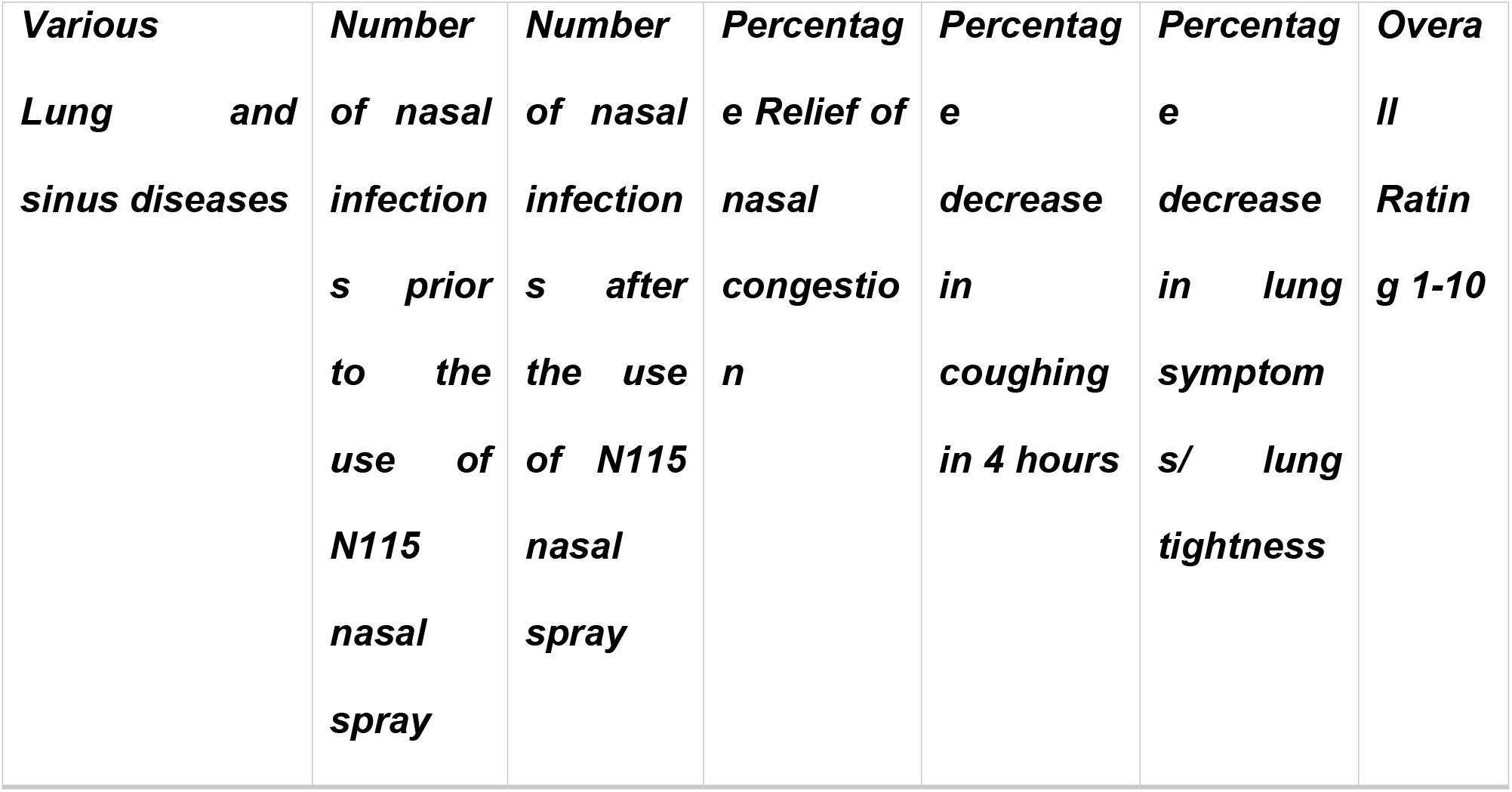
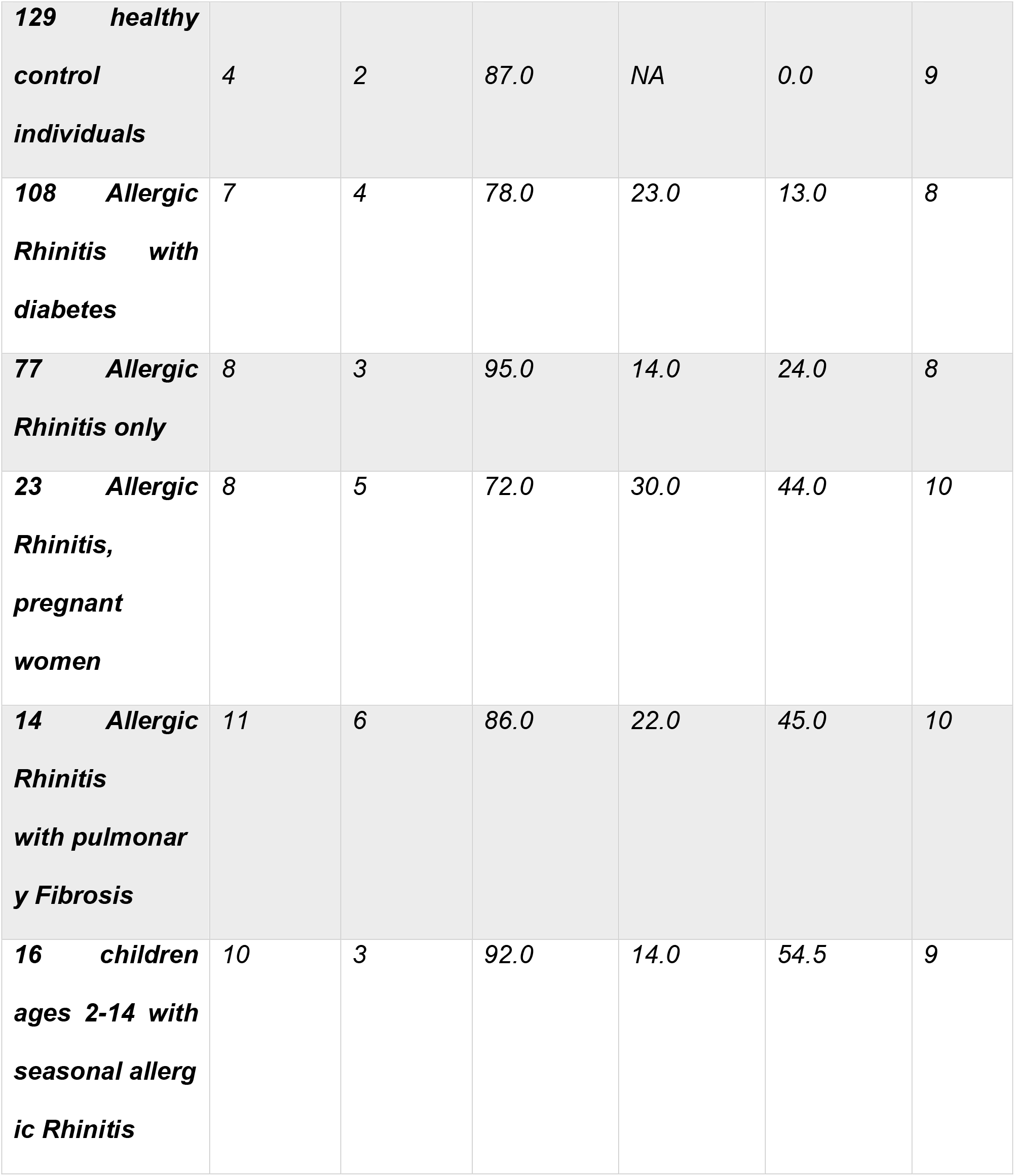
Decreasing the number, symptoms, and severity of seasonal respiratory infections. This survey of 367 patients over a two-year period shows improved number, symptoms, and severity of seasonal respiratory tract infections before and after using the 20mM sodium pyruvate nasal spray (Emphycorp, N115). Overall rating was 1-10 with 10 being the most positive result.

Based on these promising data, we sought to examine the effectiveness of NaPyr for treatment of IAV infection. We conducted preliminary toxicity experiments using nebulized NaPyr at 10mM and 1M concentrations made in house using Fisher Bioreagents NaPyr (BP356-100) diluted in PBS, or PBS for control. We found no noticeable weight loss in mice treated with 10mM NaPyr. 1M NaPyr treatment did show some slight decline in weight in mice, but this was likely due to the cloud produced by nebulizing 1M NaPyr, which was thick like chalk dust and difficult to breathe (**Figure 1A**). Overall, NaPyr was not toxic at concentrations used for treatment of IAV infected mice.

**Figure 1:**
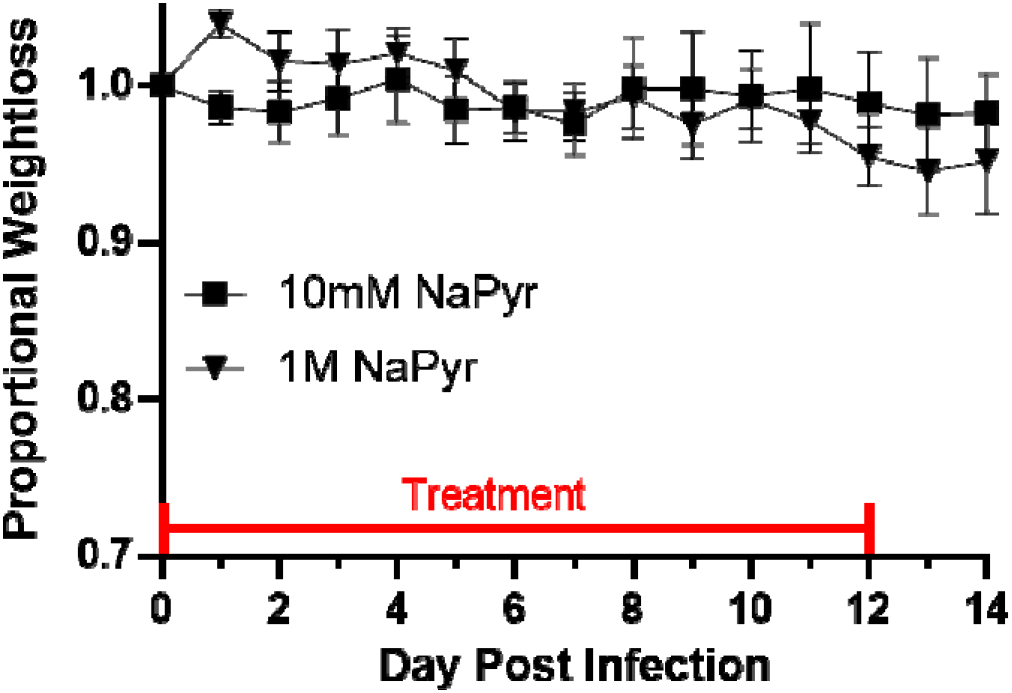
NaPyr shows no toxicity in mice. WT C57BL/6J mice were nebulized 3 times daily for 15 minutes per treatment with 10mM and 1M concentrations of NaPyr diluted in PBS for 14 days to determine toxicity and weight differences between treatment groups. Data are representative of one experiment with n=5 mice per treatment group.

### 3.2 Nebulized NaPyr improves weight loss in IAV infected mice

Our previous research with NaPyr *in vitro* established its immunomodulatory properties [23]. We, therefore, examined its effects in mice infected with IAV. WT C57BL/6J mice were infected with 250PFU of influenza A/PR/8/34 H1N1 virus and injected subcutaneously (Sub-Q) with 55mg/kg of NaPyr twice a day for 14 days and compared to PBS injection controls. Although injection of NaPyr resulted in increased food intake early and late during infection, it did not significantly improve weight loss in IAV infected mice (**Figure 2A-B**). Thus, we began looking for an alternative, more direct, administration method during infection. Aerosols are used frequently for treatment of lower respiratory infections, more specifically, viral pneumonia [27]. Hence, we hypothesized that a nebulization model would be a more direct route to the site of infection. Intriguingly, treating mice three times a day with nebulized 10mM NaPyr for approximately 15-minute intervals, resulted in some improvement in weight loss and increase in food intake (**Figure 2C-D**). (Data in Figure 2C-D are combined from 2 independent experiments with n=4-5 mice per treatment group per experiment. Two-way ANOVA p=0.0399, 0.0043, 0.0116, and 0.0363 for days 5-8 in Figure 2C, Two-way ANOVA p=0.0199 and 0.0093 for days 2-3 in Figure 2D).

**Figure 2:**
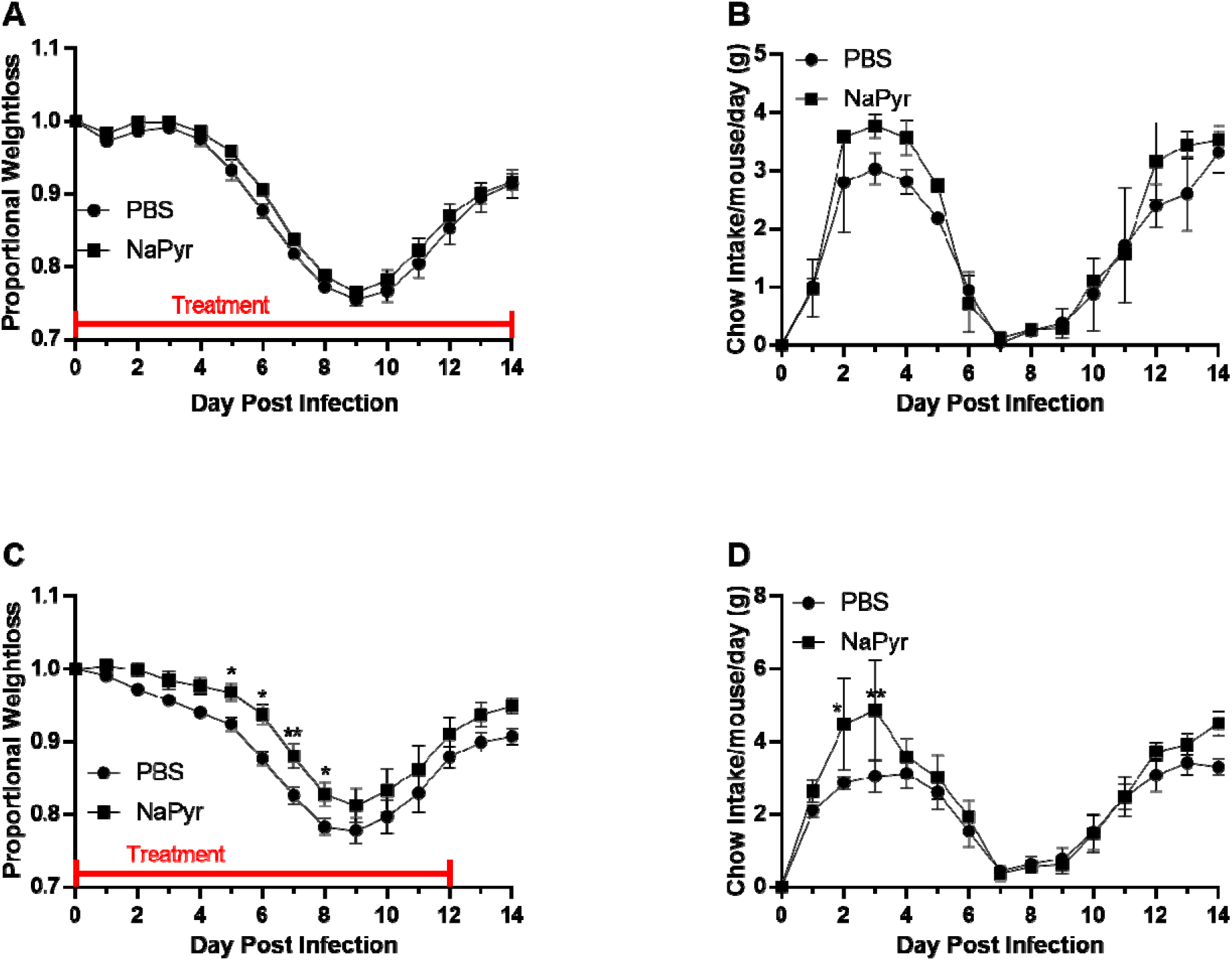
Effects of injection or nebulization of NaPyr on IAV infection. WT C57BL/6J mice were infected intranasally with 250 PFU of influenza A/PR/8/34 H1N1. Mice were treated as indicated and monitored daily for 14 days to determine survival and weight differences between treatment groups. (A) Weight loss was examined in mice injected Sub-Q with 55mg/kg NaPyr twice a day for 14-days compared to PBS injected mice. (B) Average chow intake over the 14-day IAV infection of both Sub-Q NaPyr treated and PBS treated mice. (C) Mice were treated 3 times a day with either nebulized 10mM NaPyr or nebulized PBS as control. Weight loss differences viewed over the 14-day IAV infection of both NaPyr treated and PBS treated mice. (D) Average chow intake over the 14-day IAV infection of both nebulized 10mM NaPyr and PBS treated mice. Data are representative of 2-3 individual experiments with n=4-5 mice per treatment group per independent experiment. Statistical significance was determined using a Two-way ANOVA with Fisher LSD post-hoc for multiple comparisons. * p<0.05, ** p<0.01

### 3.3 N115 decreases weight loss and increases chow intake during IAV infection

As stated above, EmphyCorp manufactures a stable 20 mM NaPyr nasal spray (N115). In Phase I/II/III clinical trials, N115 has demonstrated promising results in decreasing lung inflammation in COPD and Idiopathic Pulmonary Fibrosis patients. Using N115, we examined weight loss in WT C57BL/6J mice infected with 250PFU of influenza A/PR/8/34 H1N1 virus and treated three times a day for 20-minute intervals. Our results indicate that nebulizing mice with N115 over the course 12 days of IAV infection decreased weight loss and increased chow intake, compared to the PBS controls. We found days 7-14 to be statistically significant between the control and N115 treated groups (**Figure 3A**). Chow intake in the N115 treated mice was significantly higher on days 9-10 too (**Figure 3B**). (Weight loss and chow intake data are combined from 3 independent experiments with n=4-6 mice per treatment group per independent experiment. Two-way ANOVA p=0.0127, 0.0012, 0.0002, <0.0001, 0.0005, 0.0046, 0.0233, 0.0311 for days 7-14 respectively in Figure 3A. Two-way ANOVA p=0.0492 and 0.0335 for days 9-10 in Figure 3B).

**Figure 3:**
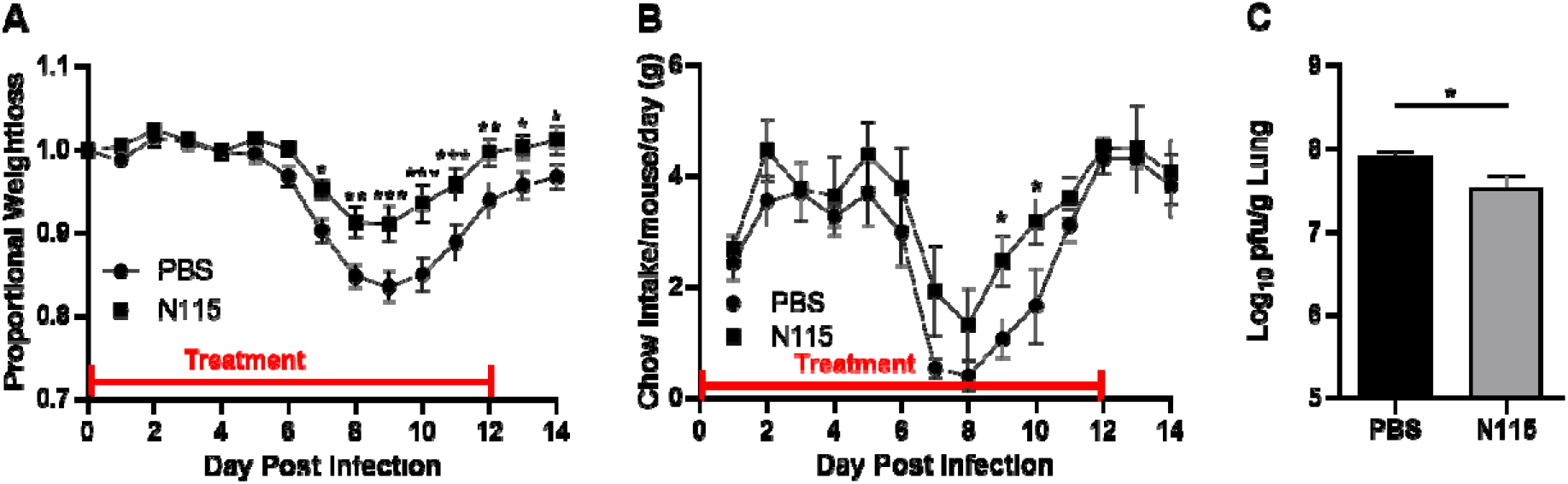
Nebulized N115 improves weight loss and virus titer in mice infected with IAV. WT C57BL/6J mice were infected intranasally with 250 PFU of influenza A/PR/8/34 H1N1. Mice were treated with either nebulized 20mM NaPyr (N115) or nebulized PBS as control 3 times a day for 20 min/treatment. (A) Mice were monitored daily for 14 days to determine weight differences between treatment groups. (B) Average chow intake over the 14-day IAV infection of both N115 treated and PBS treated mice. (C) Viral titer was assessed by plaque assay on Day 7 p.i. from lung homogenates. Weight loss and chow intake data are representative of 3 independent experiments with n=4-6 mice per treatment group per independent experiment. Viral titer data are representative of 2 individual experiments with 4-5 mice per treatment group per individual experiment. Statistical significance was determined using a Two-way ANOVA with Fisher LSD post-hoc for multiple comparisons, and Student’s T-test for single comparisons. * p<0.05, ** p<0.01, *** p<0.001

As N115 improved weight loss, we next examined the cause for improved weight loss. We investigated viral titer by plaque assay on Day 7 p.i., just as the N115 treated mice started to show improvement in weight loss. We found that there was significantly less virus in the lungs of N115 treated mice compared to PBS treated controls (**Figure 3C**). (Viral titer data are combined from 2 individual experiments with 4-5 mice per treatment group per experiment. Statistical significance was determined using a Student’s T-test, p=0.0172.)

As previously reported *in vitro* [23], we also observed significantly less IL-1β levels in the lungs of N115 treated mice (**Figure 4A**). Despite lower IL-1β levels, most leukocyte numbers were similar in the lungs of N115 and PBS treated mice, except for inflammatory monocyte numbers, which were elevated in the N115 mice (**Figure 4B-C**). Overall, N115 appears to improve disease during IAV infection by decreasing virus titers and lowering inflammatory cytokine levels. (Data in figures 4A-C are combined from 2 independent experiments with n=4-5 mice per experiment. Statistical significance was determined using a Student’s T-test. Figure 4A p=0.0351 for IL-1β; Figure 4B p=0.0442 for inflammatory monocytes).

**Figure 4:**
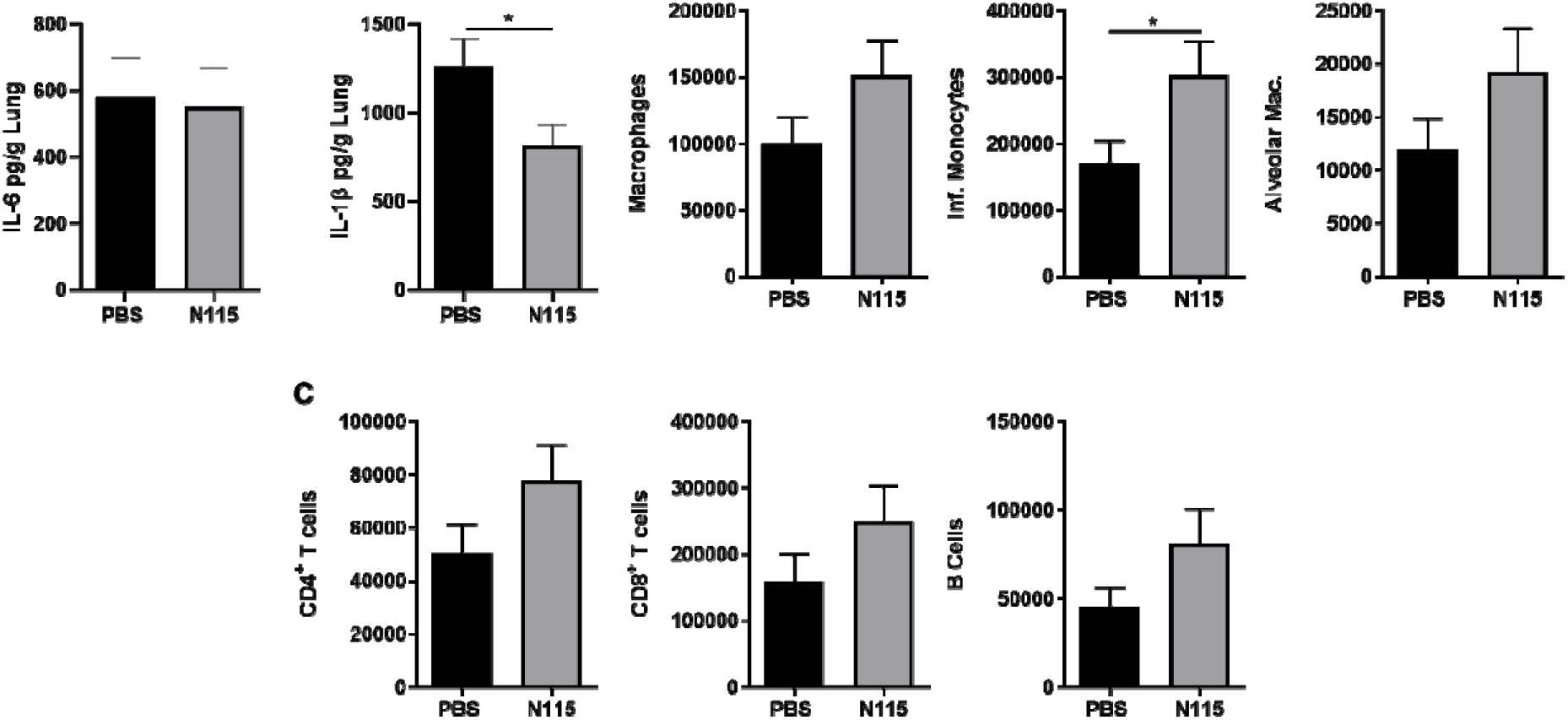
N115 treatment modulates inflammatory responses during IAV infection. WT C57BL/6J mice were anesthetized and infected with 250 PFU of influenza A/PR/8/34 H1N1. Mice were treated 3 times a daily for 20 min/treatment with either nebulized 20mM NaPyr (N115) or nebulized PBS as control. Mice were euthanized on Day 7 p.i. for tissue collection. Lung samples were then homogenized and examined via ELISA for cytokine production (A) or cellular infiltration into the lungs by flow cytometry (B-C). Data are combined from 2 independent experiments with n=4-5 mice per experiment. Statistical significance was determined using a Student’s T-test for single comparisons. * p<0.05

## 4. Discussion

Due to the evolution of antiviral or antibiotic resistance, the development of therapies that target host pathways to disrupt pathogen replication or disease is an avenue worthy of exploration. Some cellular metabolites can alter inflammation or pathogen replication, but our data suggest that the route of administrations is important. Our data indicate that sub-Q injection of NaPyr does not influence IAV induced weight loss. However, nebulizing NaPyr does have a significant impact on weight loss, virus titer and cytokine production during IAV infection *in vivo* in mice. Since Pyr can be rapidly absorbed by virtually any cell, injected NaPyr likely is taken up by other cells before reaching the target cells in the IAV infected lung [28, 29]. Most IAV antiviral treatments target specific proteins within the virus. These proteins are prone to mutations and resistance to such drugs. Certain strains of IAV are known to be resistant to the M2 and neuraminidase inhibitors [30]. Since NaPyr affects cellular metabolism and inflammation instead of directly targeting virus replication, there is a much lower chance that the virus will develop resistance to NaPyr treatment.

Influenza and COVID-19 are known to cause mortality and morbidity in the elderly and immunocompromised. However, it is often forgotten that both diseases afflict children, usually with mild symptoms. In rare cases, there is mortality caused by complications during IAV infection. Seasonally, influenza causes 7,000-26,000 hospitalizations in children under five years old [31]. COVID-19 this year has resulted in 3,240 hospitalizations in school-aged children [32]. To date, 51 children, aged less than 18, have died in the United States from complications with COVID-19 [32]. Comparatively, the CDC has reported a range of 37-188 deaths annually in children under five years old from complications caused by influenza infections [31]. Our data in this manuscript clearly demonstrate that N115 improves influenza disease. Furthermore, we have preliminary data that suggest it may work similarly during other respiratory virus infections including COVID19/SARS-CoV-2. Proactive treatments with NaPyr is not toxic and could be of benefit to children that are afflicted by many respiratory viruses.

In conclusion, we show that nebulizing mice with sodium pyruvate decreased morbidity and weight loss during infection. Additionally, treated mice consumed more chow during infection indicating improved symptoms. There were notable improvements in pro-inflammatory cytokine production (IL-1β) and lower virus titers on days 7 post infection (p.i.) in mice treated with NaPyr compared to control animals. As pyruvate acts on the host immune response and metabolic pathways and not directly on the virus, sodium pyruvate is a promising treatment option that is safe, effective, and unlikely to elicit antiviral resistance.

## Abbreviations

Pyr: Pyruvate
NaPyr: Sodium Pyruvate
IAV: influenza A virus

## 5. Author Statement

JMR and CRL performed experiments, analyzed the data and wrote the manuscript. The funders of this research had no part in the experimental design or interpretation of the data.

## 6 Acknowledgements

We would like to thank the Missouri State University Vivarium staff, especially Angela Goerndt, Shayla Lupfer, and Dr. Michael Stafford. We would also like to thank Riley Marcinczyk for help with data collection and treatments. We thank Dr. Alain Martin for his continuous communication about N115 and its current clinical findings and for financial support for this research provided by Emphycorp. We also thank Missouri State University for Graduate Student Research Funds to JMR.

